## Supplementary Figures and Tables for "Serotonin 2A receptor signaling coordinates central metabolic processes to modulate aging in response to nutrient choice": Supplementary Figures 1-9.pdf

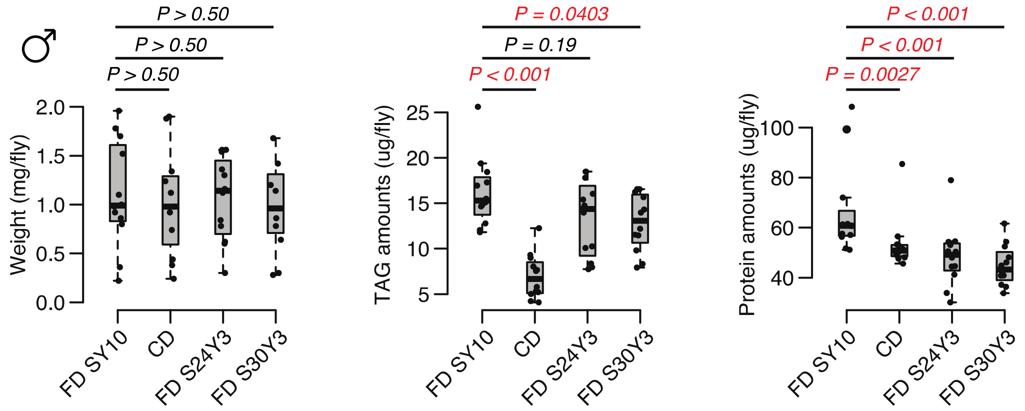


**Supplementary Figure 1**. Compared to a standard fixed diet SY10, either choice diet or high sugar diet have an effect on weight. For TAG and protein amounts, dietary choice induces a dramatic reduction on TAG amounts but only mild decrease in protein levels, whereas the effect of dietary sugar is more significant on protein amounts. N = 60 flies per each diet, which are 12 independent replicates (of 5 flies each). Comparisons were accomplished using a Mann-Whitney U test.

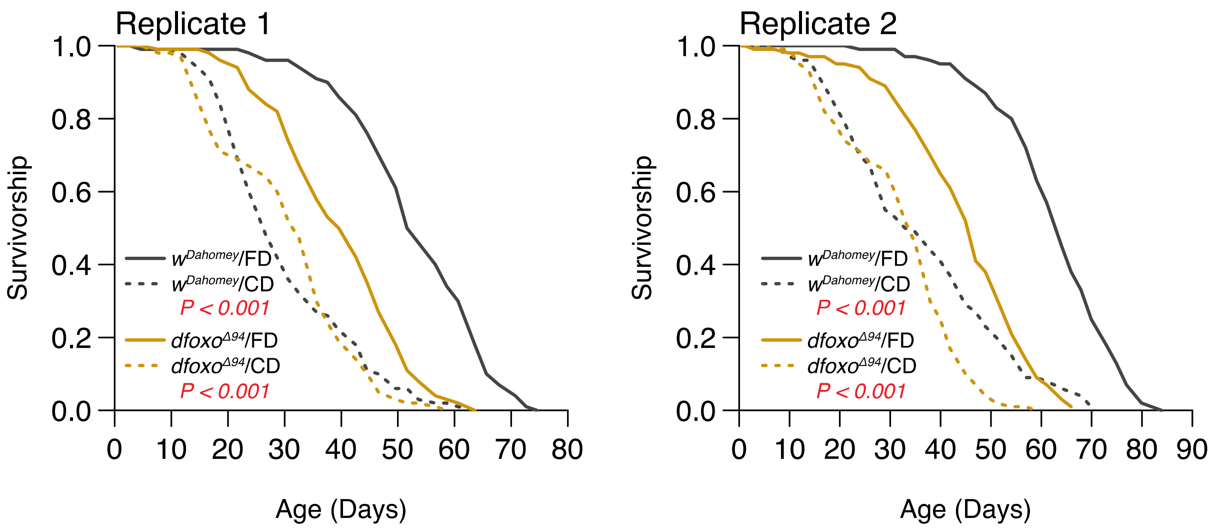


**Supplementary Figure 2.** Two independent replicates support that the lifespan effect through dietary choice is largely independent of dFOXO (N = 126-150). Comparison of survival curves was via log-rank test.

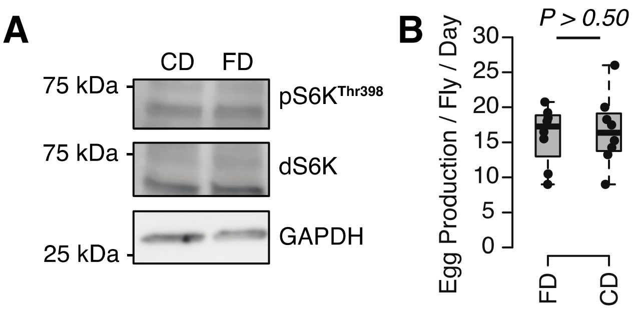


**Supplementary Figure 3**. Either TOR activity (**A**) nor egg production (**B)** responds to a choice diet. For western blot, 10 flies are pooled for each sample. For egging, 8 biological replicates were used per diet (of 8 females and 8 males each). Measures were taken over four days (see our methods). *P*-value was obtained using a Mann-Whitney U test.

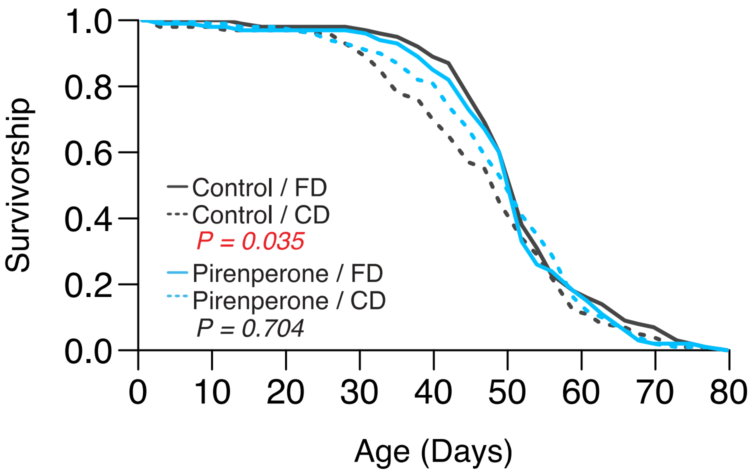


**Supplementary Figure 4**. Second replicate to validate the role of pirenperone in abrogating the lifespan effect of dietary choice. *P*-values were obtained using a log-rank test (N = 130-144).

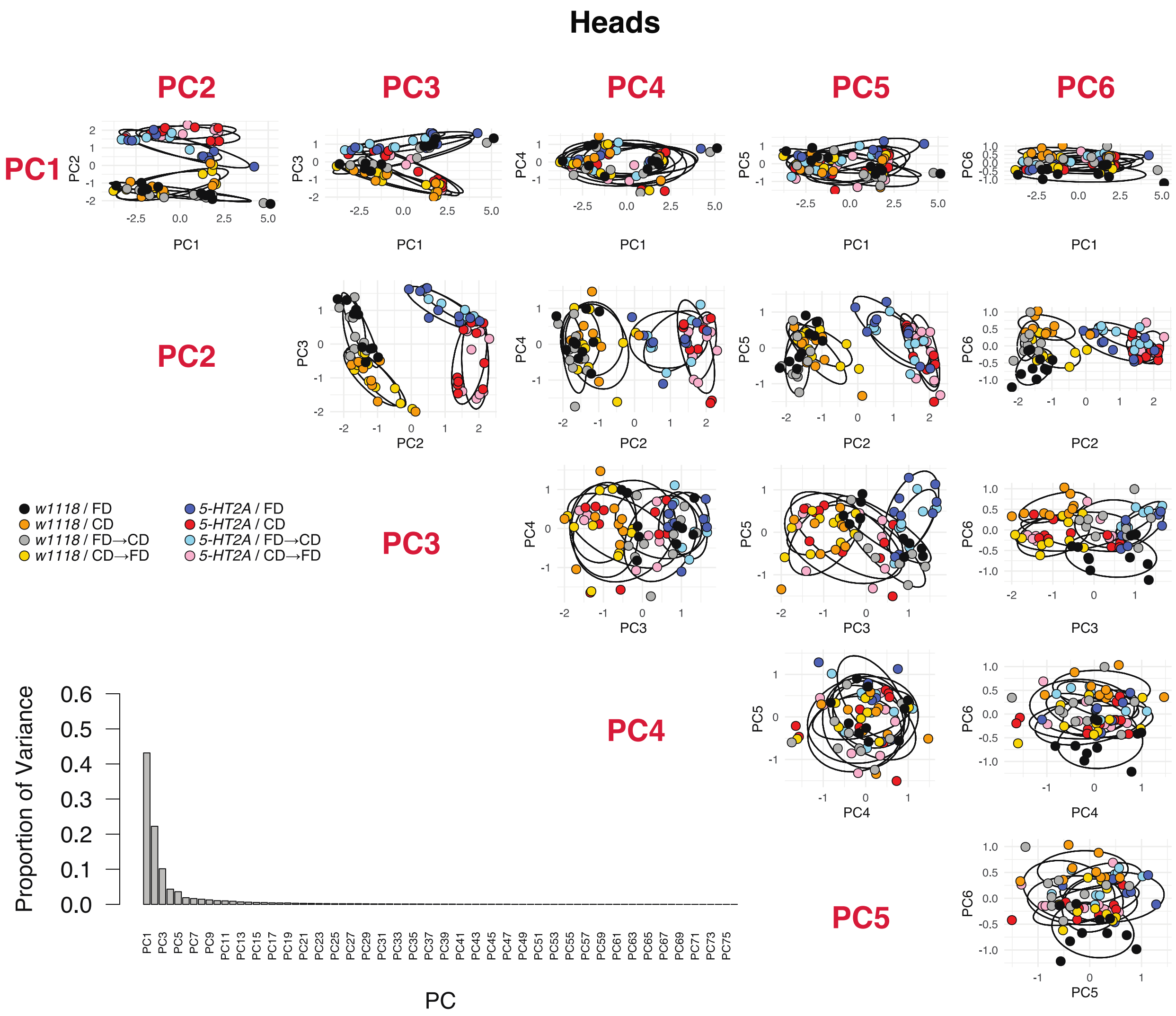


**Supplementary Figure 5**. PCA analyses in head metabolomes, including the proportion of variance of each PC and the PCA plots for the first six PCs.

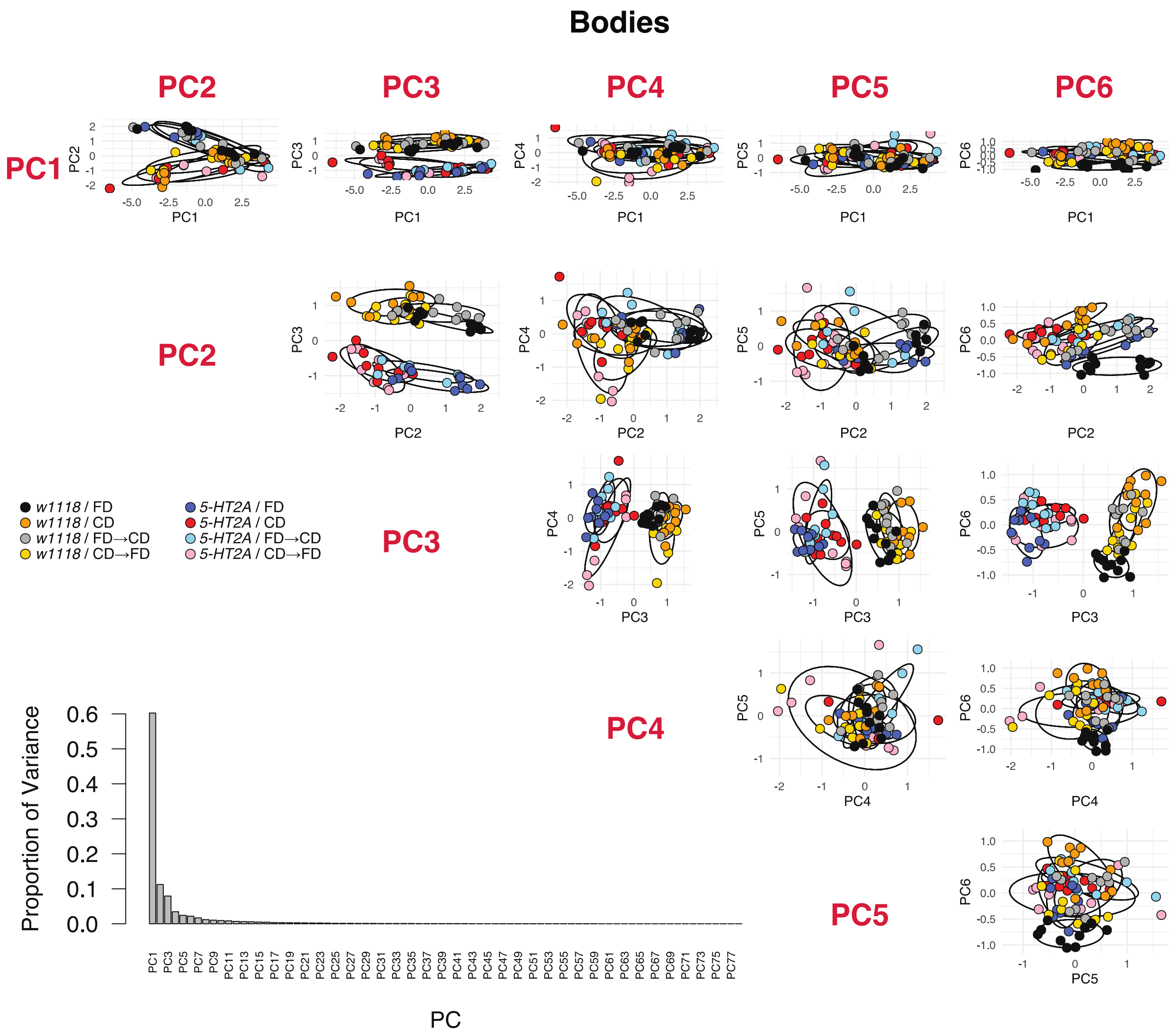


**Supplementary Figure 6**. PCA analyses in body metabolomes, including the proportion of variance of each PC and the PCA plots for the first six PCs.

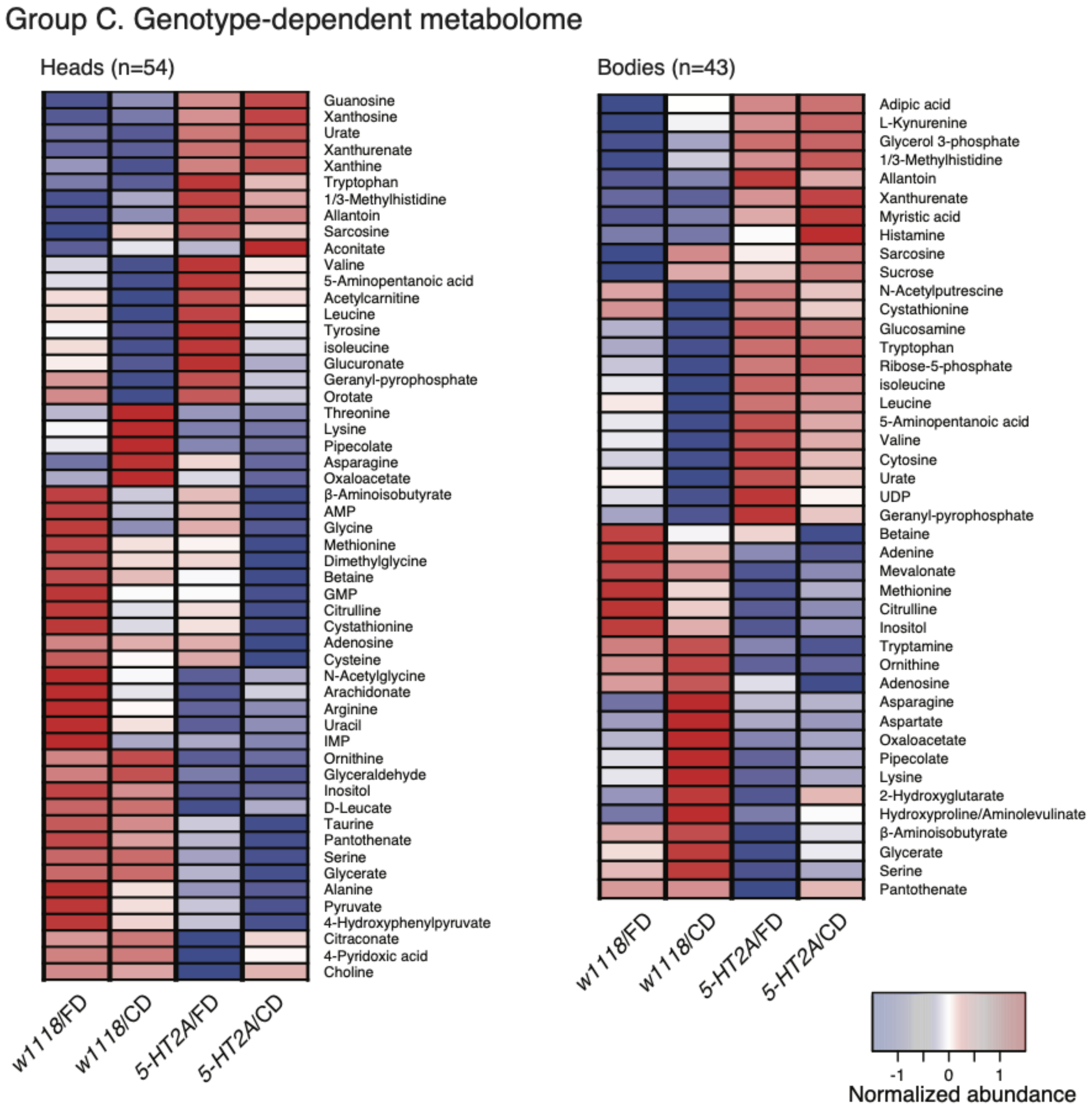


**Supplementary Figure 7**. Clustering of the abundance of each metabolite that differed between genotypes in head or body metabolomes.

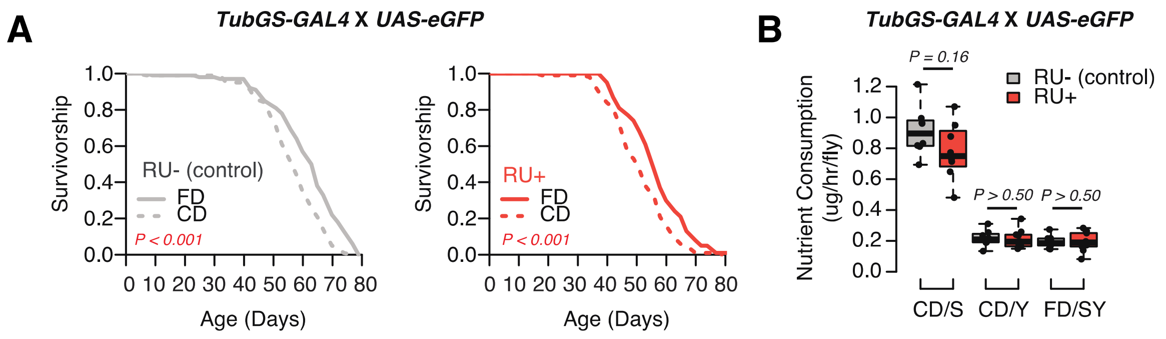


**Supplementary Figure 8.** (**A**) GeneSwitch control flies (*TubGS-GAL4* X *UAS-eGFP*) still exhibit shortened lifespan on a choice diet, in both RU+ and RU- conditions, indicating RU486 has no impact on the choice effect. N = 143-152. Comparison of survival curves was via a log-rank test. (**B**) Similarly, RU486 has no effects in nutrient consumption on either diet (N=60-79, *P*-values were obtained using a Mann-Whitney U test).

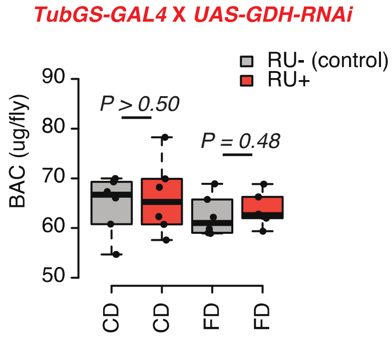


**Supplementary Figure 9.** Knock-down *GDH* has no effects on the protein amount of flies. N=60, including six independent samples per each group (of 10 flies each). *P*-values were obtained using a Mann-Whitney U test.
