## Supplementary Figures and Tables for "Serotonin 2A receptor signaling coordinates central metabolic processes to modulate aging in response to nutrient choice": Supplementary Tables 1-2.pdf

**Supplementary Table 1**. Summary of dietary choice effects on lifespan of flies from different wildtype strains. Values represent mean lifespan (SEM) in days for each cohort. *P*-values are from log-rank ratio test.

| Sex | Strain | Fixed Diet | Choice Diet | %Change | *P*-value |
|---|---|---|---|---|---|
| Male | *Canton-S* | 60.85 (1.48) | 47.23 (1.55) | 28.84% | < 0.001 |
| | $w^{1118}$ Pletcher lab strain1 | 59.73 (1.48) | 46.19 (1.65) | 29.31% | < 0.001 |
| | $w^{1118}$ Pletcher lab strain2 | 61.59 (1.90) | 44.86 (1.96) | 37.29% | < 0.001 |
| Female | *Canton-S* | 54.71 (0.96) | 52.02 (0.96) | 5.17% | 0.033 |
| | $w^{1118}$ Pletcher lab strain1 | 54.56 (1.04) | 51.21 (1.29) | 6.54% | 0.246 |
| | $w^{1118}$ Pletcher lab strain2 | 52.48 (0.90) | 45.65 (1.25) | 14.96% | 0.010 |

**Supplementary Table 2**. Summary of 5-HT2A gene expression in various tissues. RNA-seq data are obtained from FlyAtlas2 (flyatlas.gla.ac.uk/FlyAtlas2/).

| | Adult Male | | Adult Female | |
|---|---|---|---|---|
| **Tissue** | **FPKM** | **Enrichment** | **FPKM** | **Enrichment** |
| Head | 23 | 4.6 | 17 | 8.7 |
| Eye | 95 | 19 | 72 | 36 |
| Brain / CNS | 16 | 3.1 | 17 | 8.4 |
| Thoracicoabdominal ganglion | 7.9 | 1.6 | 9.3 | 4.7 |
| Crop | 3.3 | 0.7 | 5.7 | 2.9 |
| Midgut | 0.8 | 0.2 | 0.2 | N.A. |
| Hindgut | 0.9 | 0.2 | 0.4 | N.A. |
| Malpighian Tubules | 0.3 | 0.1 | 0.3 | N.A. |
| Salivary gland | 172 | 34 | 138 | 69 |
| Ovary | | | 0.1 | N.A. |
| Virgin Spermatheca | | | 0.3 | N.A. |
| Mated Spermatheca | | | 0.1 | N.A. |
| Testis | 5.0 | 1.0 | | |
| Accessory glands | 8.5 | 1.7 | | |
| Carcass | 3.1 | 0.6 | 3.2 | 1.6 |
| Rectal pad | 2.2 | 0.4 | 2.2 | 1.1 |
